# The dynamics of coexistence and succession in a decaying ecosystem

**DOI:** 10.1101/2025.04.28.650976

**Authors:** Ata Kalirad, Penghieng Theam, Ralf J. Sommer

## Abstract

Elucidating the rules that govern community assembly and enable the coexistence of species is central to ecology. Much of the current understanding revolves around the composition of communities at equilibrium. In contrast, transient ecological dynamics, in particular the succession of species following an environmental disturbance, remain largely unexplored. We present *Gymnogaster buphthalma* beetle carcasses as a model to study species coexistence in disequilibrium. In this community, nematodes appear in succession – with multiple feeding and reproductive strategies, transience, and associated dispersal – creating a perfect model for metacommunity ecology. We computationally reconstructed decaying *G. buphthalma* beetles sampled from the wild using experimentally-derived life history data, i.e. emergence times, sex ratios, and fecundity measurements. Through agent-based modeling, we show that nematode coexistence is possible over a wide range of scenarios. This work provides a unique experimental and computational framework to synthesize various approaches to metacommunity and species succession theories.

## Introduction

The importance of transient dynamics in shaping short-lived ecosystems is increasingly being recognized (*1–6*). Particularly, the rapid transformation of habitats due to global climate change (*7, 8*) necessitates experimental and theoretical work to better understand the ecological dynamics in transient ecosystems. Filling this knowledge gap would also relate the paradox of coexistence in ecological communities to the long-standing problem of species succession, which plays a major role in shaping the composition of short-lived communities (*9, 10*). This paradox – the observation that, instead of competitive exclusion, species can coexist in a community – remains a central problem in ecology (*11, 12*). Mechanistic solutions postulate long-term coexistence as an asymptotic state resulting from various forms of competitions due to limited resources, predation, and other ecological interactions (*13*). In contrast, the neutral approach interprets the observed coexistence as a short-lived state, shaped by chance, before an inevitable monodominance (*14, 15*).

Deterministic models predict long-term coexistence over time scales approaching infinity in communities where species densities can vary in a continuous fashion (*16*). By their very design, such models exclude the consequences of discrete individuals in a population, e.g., demographic stochasticity (*17, 18*). Therefore, the patterns of coexistence in ecosystems with rapid turn-over, specifically those that follow a boom-and-bust dynamic, remain largely unexplored. Despite extensive studies of plant communities, little is known about transient coexistence during succession. In invertebrates, one notable study system is decaying insect carcasses frequently found in the soil, which are characterized by their organismal diversity and rapid decomposition. As such, decaying insects are a perfect model to study succession and coexistence over short periods of time (*19, 20*).

Here we take advantage of *Gymnogaster buphthalma* beetles of the Scarabaeidae family as a model to study succession and coexistence in a decaying ecosystem. Various free-living nematodes are found in association with these and other scarab beetles (*21*). We integrate laboratory data of nematode species that emerged from *G. buphthalma* beetles sampled on Réunion island with an agent-based modeling approach to understand the dynamics of species coexistence and succession.

## Results

### The community composition emerging on *G. buphthalma* beetle carcasses

For this study, *G. buphthalma* beetles sampled from Réunion island were sacrificed in the laboratory on nematode growth medium (NGM) plates (Fig. 1A). As long as beetles are alive, nematodes stay in the arrested dauer stage. Following the beetlés death, microbes decompose the carcass, providing food for nematodes, which in response exit the dauer stage and develop into adults. Following the sacrifice, plates were regularly observed to accurately infer nematode emergence time on the carcasses (Fig. 1B). All 24 *G. buphthalma* beetles harbored nematodes with three distinct patterns. Interestingly, all beetles contained isolates of a previously unknown *Acrostichus* species, which will be described elsewhere. On 13 carcasses, the presence of this species were accompanied by isolates of *Pristionchus pacificus* – an established model organism (*22*) – or *Pristionchus mayeri* (*23*). Importantly, isolates of *P. pacificus, P. mayeri*, and *Acrostichus* sp. emerged together on 6 beetles (Fig. 1B). In addition to observing the nematode composition patterns, we were able to accurately estimate the timing of emergence of each nematode on a given beetle. This allowed us to incorporate differences in the emergence strategies in addition to life history measurements into an integrative modeling framework to reconstruct the dynamics on a decaying beetle.

**Figure 1.**
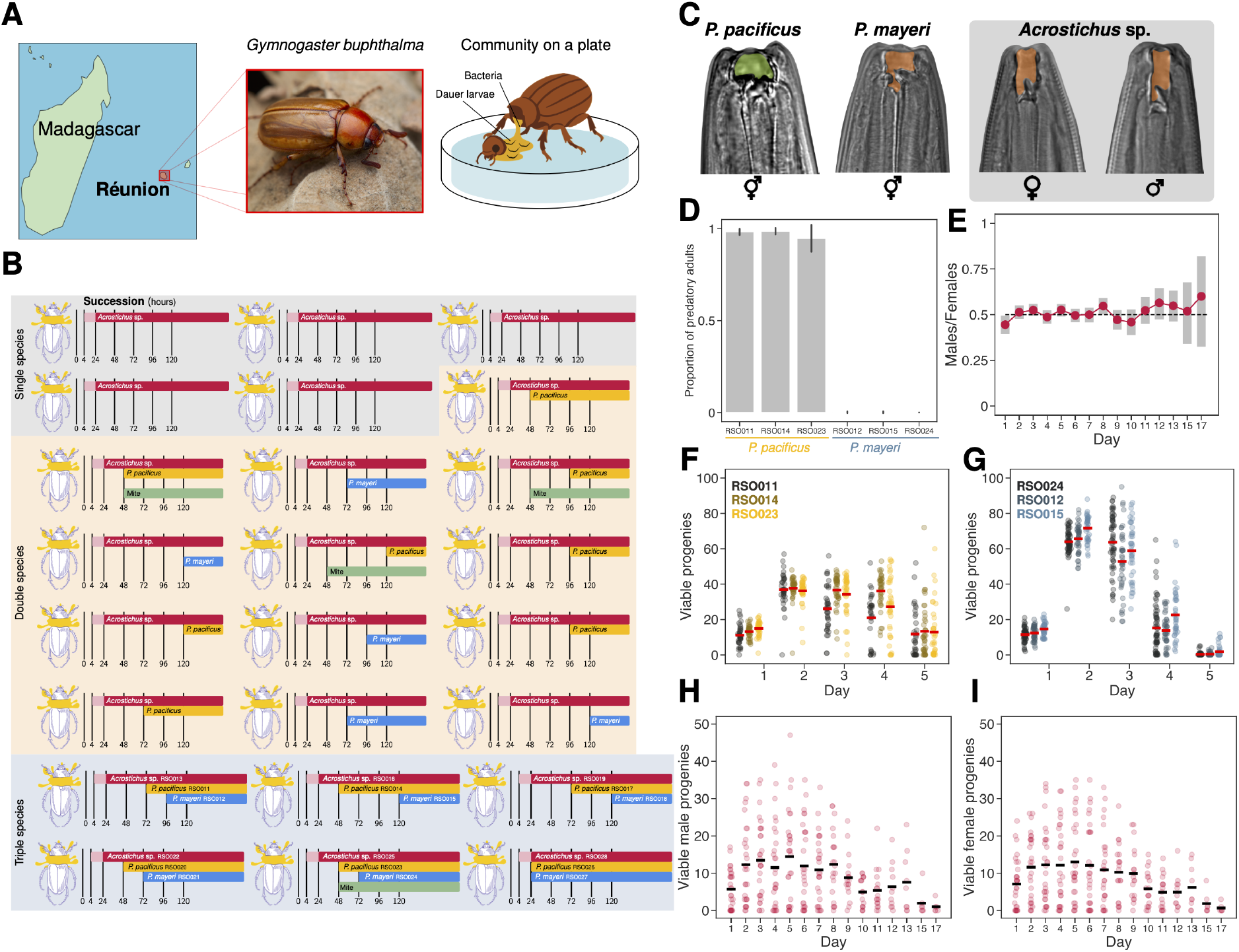
Emergence dynamics and composition of nematode communities on beetle carcasses. (A) *G. buphthalma* beetles sampled from Réunion island were used to construct “community on a plate” (Photo of *G. buphthalma* courtesy of Zeeshan Mirza). (B) Two or more nematode species emerged from the majority of *G. buphthalma* carcasses (19 out of 24), with six containing various isolates belonging to three different nematode species. (C) The three-species communities represent diverse feeding structures: the *P. pacificus* isolates develop the wide-mouth eurystomatous mouth form, which enables killing of competing nematodes, while the *P. mayeri* and *Acrostichus* sp. isolates develop feeding structures suited to bacterivory. Additionally, both *Pristionchus* species are androdioecious hermaphrodites, in contrast to *Acrostichus* sp., which is a gonochoristic species. (D) The proportion of predatory adults in the isolates of *P. pacificus* and *P. mayeri*. (E) *Acrostichus* sp. RSO013 progeny over the entire egg-laying period showed a 1:1 sex ratio. Vertical bars represent the 95% the highest density interval (HDI) for the sex ratio based on a binomial model. (F) Fecundity of the *P. pacificus* isolates. (G) Fecundity of the *P. mayeri* isolates. The number of viable male (H) and female (I) progenies of *Acrostichus* RSO013. Horizontal lines in (F) to (I) represent the mean values.

The communities containing three nematode species provide a rare opportunity to investigate community dynamics of a transient ecosystem with three unique characteristics, i) a diversity of feeding structures, ii) different reproductive strategies, and iii) a boom-and-bust life cycle (Fig. 1C) (*24*). *P. pacificus* isolated from the beetles developed the wide-mouthed (eurystomatous) form with two teeth, which enables them to kill juveniles of other nematodes in addition to consuming bacteria (Fig. 1D) (*25, 26*). In contrast, *P. mayeri* and *Acrostichus* sp. developed bacteriovorus feeding structures under the same condition (Fig. 1D). Thus, the beetle carcass is inhabited by nematodes with different feeding strategies including life history intraguild predation by *P. pacificus* (*27*). However, given the amount of surplus killing observed in the lab, this interaction can also be described as interference competition (*28*).

Intriguingly, these nematodes also reflect a plethora of reproductive strategies (Fig. 1C). Both *Pristionchus* species encountered on these *G. buphthalma* cadavers are androdioecious hermaphrodites that can only reproduce by selfing because males were not found among the emerging nematodes. In contrast, *Acrostichus* sp. is a gonochoristic species with distinct males and females, indicating that beetle carcasses are populated by free-living nematodes with vastly different reproductive strategies. Therefore, with respect to reproductive strategies, the presence of these three species raises a fascinating question: Given the well-known cost of males, which has been hypothesized to handicap gonochorist species in competition with hermaphrodites (*29, 30*), how could *Acrostichus* sp. compete with its two rivals? Additionally, the cost of males in these competitions will be confounded with intraguild predation by *P. pacificus*.

We attempted to reconstruct the dynamics of coexistence and succession using an approach that combines laboratory measurements of nematode life history traits with an agent-based modeling framework (Fig. 2). Firstly, we measured the sex ratio of the *Acrostichus* sp. RSO013 under laboratory conditions, which indicated an equal proportion of males to females over life-time fecundity (Fig. 1E). Secondly, we measured the fecundity of the three nematode species (Fig. 1F–I). For that, we quantified the number of progenies produced by a single mother during a 24-hour time period for a total of 7 days. The fecundities were consistent amongst the isolates of *P. pacificus* and *P. mayeri*, with the isolates of the latter species having consistently higher fecundities relative to the former (Fig. 1F–G, Fig. S1). For the gonochorist *Acrostichus* sp., we followed mating pairs over 17 days. Total progeny number is much higher for *Acrostichus* relative to the hermaphroditic *Pristionchus* species (Fig. S1). This observation reflects the role of sperm availability as the limiting factor in nematode life-time fecundity. Taken together, these measurements provide a treasure trove of data, enabling a computationally-thorough reconstruction of the dynamics in these communities.

**Figure 2.**
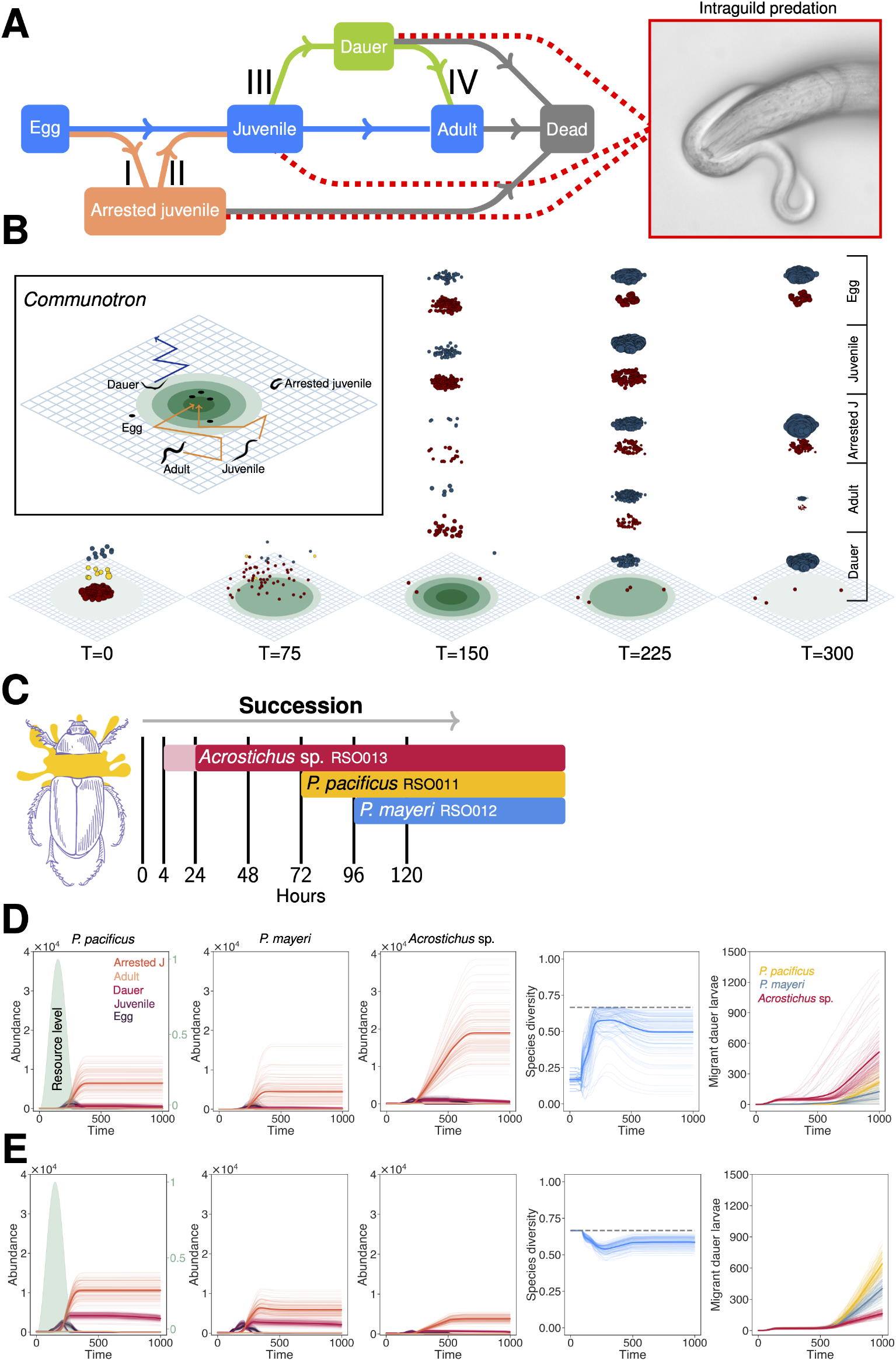
Computational reconstruction of nematode communities using *Communotron*. (A) Nematodes in *Communotron* follow a realistic life cycle, which includes alternative developmental states, namely, arrested juvenile and the dauer state. Transitions I to IV represent alternative developmental paths which are determined by local population density and/or resource availability. *P. pacificus* adults can kill juvenile, dauer, and arrested juvenile stages of *P. mayeri* and *Acrostichus* sp., resulting in life-history intraguild predation. (B) Worms in *Communotron* occupy points on a two-dimensional lattice. Bacterial growth is simulated as a dynamic component of the model, generating a time-dependent gradient that affects the bias of the movement of the motile stages in the community and the probabilities of transitioning to alternative developmental states. (C) The pattern of succession on a *G. buphthalma* beetle as observed on a plate. We used *Communotron* to computationally reconstruct the dynamics of this community based on laboratory measurements and the succession timeline. (D) The community dynamics based on 100 simulations of *Communotron* (C). The species diversity is calculated using the Gini–Simpson index, with the dotted gray line indicating the maximum expected value, 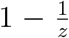, where *z* is the number of species. (E) Simulation of the community under the counterfactual condition of equal starting population sizes for the three species. Parameters used in these simulations: initial number of dauers: 200 *Acrostichus* sp., 10 *P. mayeri*, 10 *P. pacificus*; *m* = 100 *×* 100, *η* = 0.005, *L* = 10; in the counterfactual scenario, all the parameters are the same, except for starting population sizes: 200 *Acrostichus* sp., 200 *P. mayeri*, 200 *P. pacificus*.

### *Communotron*: an agent-based reconstruction of a decaying beetle

To computationally reconstruct the dynamics of competition between nematodes on decaying *G. buphthalma* beetles, we constructed an agent-based model, referred to as *Communotron* (Figs. 2A–2B). This model is inspired by a model previously introduced to illustrate the role of phenotypic plasticity in competition among nematodes (*31*). In *Communotron*, individual worms follow a realistic life cycle, which includes environmentally-dependent transitions to alternative developmental paths – i.e., arrested juvenile and dauer stages – and include life-history intraguild predation by *P. pacificus* adults (Fig. 2A). In short, *Communotron* enables an agent-based framework that combines the ability to include experimental data, resource dynamics, and simulation on the scale of hours (For more detail, see Materials and Methods).

### The cost of males in *Acrostichus* sp. is mitigated by its dauer-exit strategy

To illustrate the workflow of *Communotron*, we reconstructed the community observed on one of the beetles, which contained *Acrostichus* sp. RSO013, *P. pacificus* RSO011, and *P. mayeri* RSO012 (Fig. 2C). In this community, adult *Acrostichus* sp. emerged 24 hours after the beetle was sacrificed, whereas *P. pacificus* and *P. mayeri* emerged after 48 and 72 hours, respectively (Fig. 2C). The reconstructed communities indicate that the early emergence of *Acrostichus* sp. mitigates much of its shortcomings as a competitor, resulting in communities where this species composes most of the biomass at every developmental stage. In addition, given that dauer larvae can exit the local community, this strategy results in much higher migrant dauer larvae belonging to *Acrostichus* sp. compared to its competitors (Fig. 2D).

It is notable that on all 24 observed communities, the *Acrostichus* sp. isolates emerged early out of their dauer stage, even in the absence of any other nematode on the decaying beetle (Fig. 1B). Thus, we hypothesized that the dauer-exist strategy in *Acrostichus* sp. – which ensures a head start relative to hermaphroditic and potentially predatory competitors at the expense of having access to higher levels of resource during these initial decomposition phase – can offset both the cost of having males in this species and its non-predatory feeding structure. The results of *Communotron* are in line with this hypothesis.

The observed dynamics is strongly affected by the number of dauer larvae of each species present at the start of the formation of the community. To illustrate this point, we simulated the same community under the counterfactual condition in which all of the three species emerged on the beetle carcass in equal numbers. Under this condition, *Acrostichus* sp. performs worse than its competitors. However, none of the species are competitively excluded from the community and dauer larvae of all three species can be found in the surrounding environment (Fig. 2E). Taken together, these findings illustrate how the theoretical cost of males can be mitigated in nature.

### Species coexist during pairwise competition on the beetle carcass

The three-way interaction between these nematode species was observed in 25% of the analyzed beetles, whereas more than 50% carry *Acrostichus* sp. together with one of the *Pristionchus* species. These two-way interactions differ from each other because *Acrostichus* sp. competes with a potential predator only when it co-occurs with *P. pacificus*. To investigate how coexistence is possible in these two-way interactions, we simulated communities on decaying beetles that contain a pair of species (Fig. 3).

**Figure 3.**
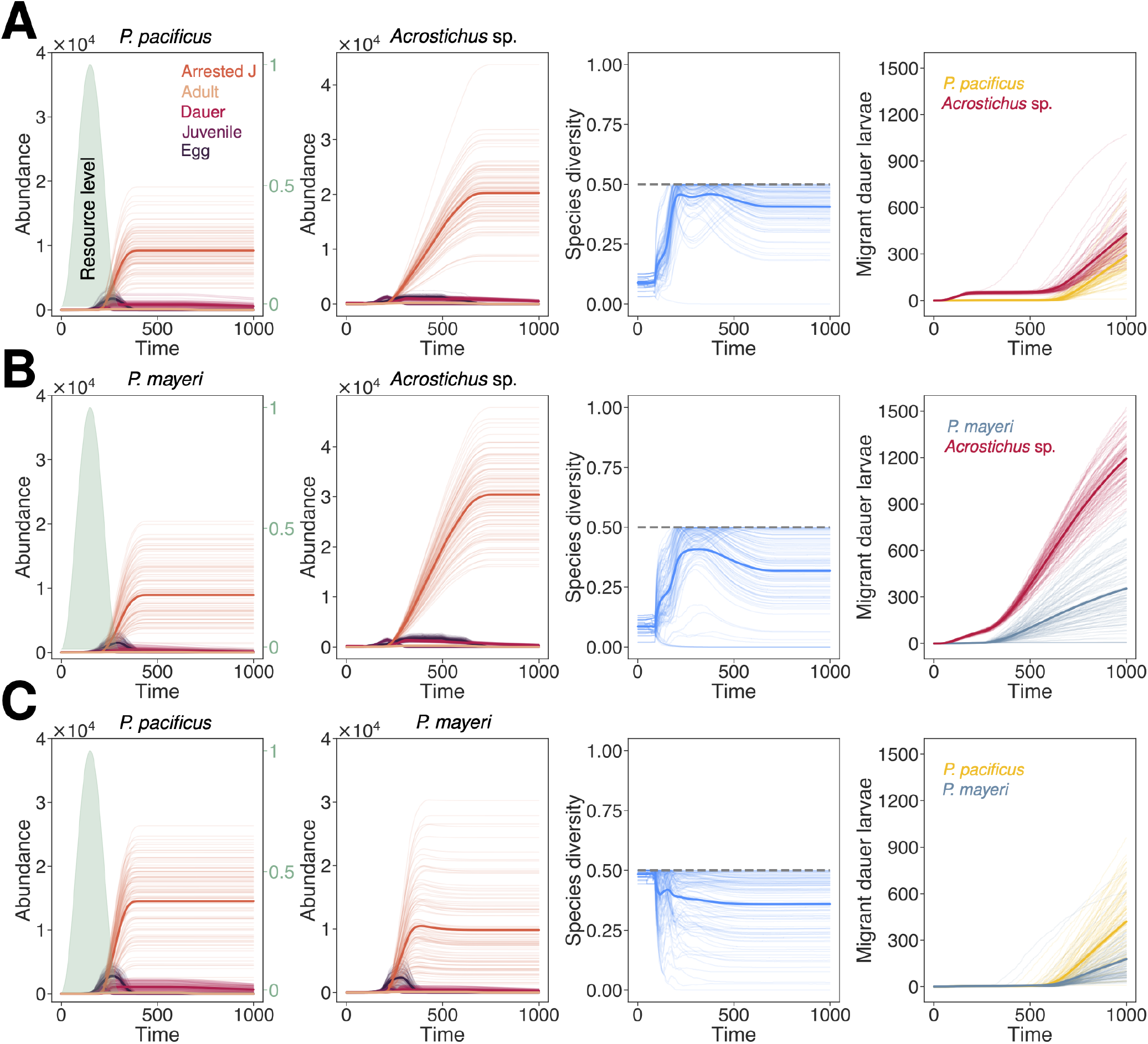
Coexistence during pairwise competition. (A) Pairwise competition between *P. pacificus* and *Acrostichus* sp., (B) pairwise competition between *P. mayeri* and *Acrostichus* sp., and (C) pairwise competition between *P. mayeri* and *P. pacificus*. Parameters used in these simulations: initial number of dauers: 200 *Acrostichus* sp., 10 *P. mayeri*, 10 *P. pacificus*; *m* = 100 *×* 100, *η* = 0.005, *L* = 10.

Pairwise competition between *P. pacificus* and *Acrostichus* sp. should be both affected by the cost of males in *Acrostichus* sp. and the effect of intraguild predation exerted by *P. pacificus*. It is possible *a priori* that the combined effect would result in the competitive exclusion of the gonochorist population on the carcass. However, we found that these species can coexist as long as resources are available (Fig. 3A). In comparison, pairwise competition between *P. mayeri* and *Acrostichus* sp. is intriguing since it is directly a result of cost of males in *Acrostichus* sp. versus a hermaphroditic competitor, who is bacteriovorus and does not engage in predation. Interestingly, these two species can also coexist in a community (Fig. 3B). It should be noted that the realized cost of males is both affected by the dauer-exit strategy of *Acrostichus* sp. and the initial number of dauer larvae of each species on the beetle (Fig. S2). Thus, similarly as in the three-species communities explored before, multiple factors can overcome the cost of males at the community level.

Finally, our modeling approach also allows us to investigate the interaction of the two *Pristionchus* species in isolation even though this is not observed in nature (Fig. 3C). Hypothetically, the pairwise competition between *P. pacificus* and *P. mayeri* should be the net outcome of the effect of intraguild predation by *P. pacificus* and the higher fecundities of *P. mayeri*. Does the interplay between factors prevent coexistence as long as resource is available? The results of our simulations indicate that, while intraguild predation reduces the proportion of *P. mayeri* in the community, both species still coexist (Fig. 3C). Thus, coexistence is observed across a wide range of conditions.

### Longer resource availability does not hinder coexistence

The outcome of competition on a decaying beetle differs from a stochastic model of population dynamics without resource dynamics, which predicts all-or-nothing outcomes (*31*). One could imagine that the transient coexistence in our model is purely a function of the single phase of resource availability, and would vanish if bacterial load was periodically replenished on the carcass. Given that a decaying beetle carcass in the soil is an open system, and other microbes can enter it, it is not unrealistic to imagine multiple phases of resource availability. To test this notion, we first simulated communities of three species with two phases of bacterial growth (Fig. 4A). Interestingly, the addition of a second resource availability phase resulted in communities in which all three species still coexist. But even more surprisingly, under the same conditions, a counterfactual scenario in which the dauer larvae of all three species emerge from the carcass in equal quantities promotes coexistence (Fig. 4B). This result differs from similar simulations with only a single resource availability phase. Thus, the periodic availability of resource promotes coexistence, instead of hindering it. This point can be made even more explicit if we simulate an extreme scenario, in which bacterial population is perpetually replenished. Here, the coexistence of all three species is still observed, with *Acrostichus* sp. RSO013 accounting for the majority of dauer larvae produced on a carcass (Fig. 4C).

**Figure 4.**
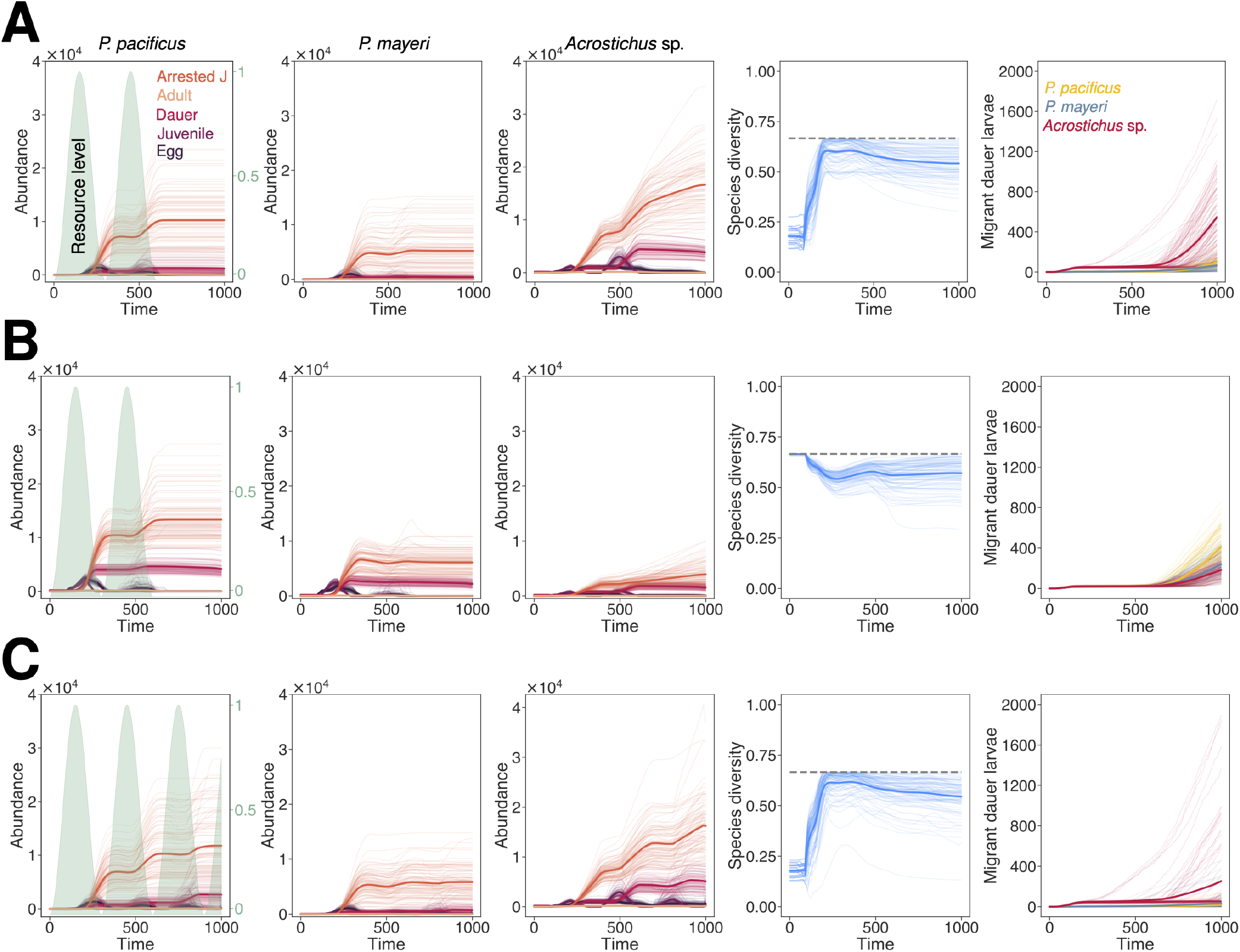
Counterfactual scenarios of resource availability and coexistence. (A) Three-way competition between *P. pacificus, P. mayeri*, and *Acrostichus* sp. with two phases of resource availability. (B) Another counterfactual scenario in which all three species exit the dauer stage on a decaying carcass in equal numbers with conditions identical to (A). (C) Three-way competition between *P. pacificus, P. mayeri*, and *Acrostichus* sp. with multiple phases of resource availability. Parameters used in these simulations: initial number of dauers in (A) and (C): 200 *Acrostichus* sp., 10 *P. mayeri*, 10 *P. pacificus*; *m* = 100 *×* 100, *η* = 0.005, *L* = 10.

Taken together, the communities reconstructed using *Communotron* with single or multiple resource availability phases illustrate how a boom-and-bust dynamic results in competitive outcomes that are different from a model without explicit resource availability dynamic. While the latter predicts competitive exclusion, the former indicate conditions under which species can coexist, in spite of *a priori* disadvantages attributed to each competitor in isolation.

## Discussion

Much of our understanding of community ecology revolves around the composition of communities at equilibrium and their stability following perturbations (*32–34*). However, the importance of disequilibrium in shaping ecological dynamics has been noted since the inception of spatial ecology, further highlighted by the introduction of patch dynamics, which in turn led to the establishment of metacommunity ecology (*35–37*). Much of the theoretical contributions in ecology remains rooted in classical models designed to explore ecosystems at equilibrium. Furthermore, incommensurable predictions derived from various metacommunity theories — the patch dynamics model, the neutral theory model, the species sorting model, and the mass effect model, each emphasizing distinct processes — have prompted an ongoing research program to provide an integrative “Übermodel” that could integrate all these processes (*37*).

The importance of the combined effects of various ecological processes at disequilibrium is probably nowhere as prominent as when multiple species occupy an emergent ecosystem following an environmental disturbance. This phenomenon of species succession (*38,39*) has been extensively documented, albeit almost exclusively in plants (*40*). In this context, the decaying *G. buphthalma* beetle provides a comprehensive model for metacommunity ecology, since it incorporates various processes relevant to metacommunity theories, e.g., dispersal, predation, demographic heterogeneity, and reproductive modes (Fig. 5). These processes are intertwined during the formation of a habitat following a disturbance, i.e., the natural death of the beetle. The community on the dacaying carcass is established with a limited number of dauer larvae. During the initial colonization phase, stochastic events can lead to ecological drift, which has long been suggested to play a crucial role in shaping community composition when the initial community size is small (*41–44*). This ecosystem lasts as long as microbial resources remain available, a case of species succession in disequilibrium. Our computational reconstruction of the decaying beetle indicates, contrary to many deterministic models, the breath of scenarios that promote transient coexistence in such a model. This framework can be used to study the competition-colonization trade-off models to better understand the relative importance of factors such as local competitiveness and dispersal in a natural system (*45, 46*).

**Figure 5.**
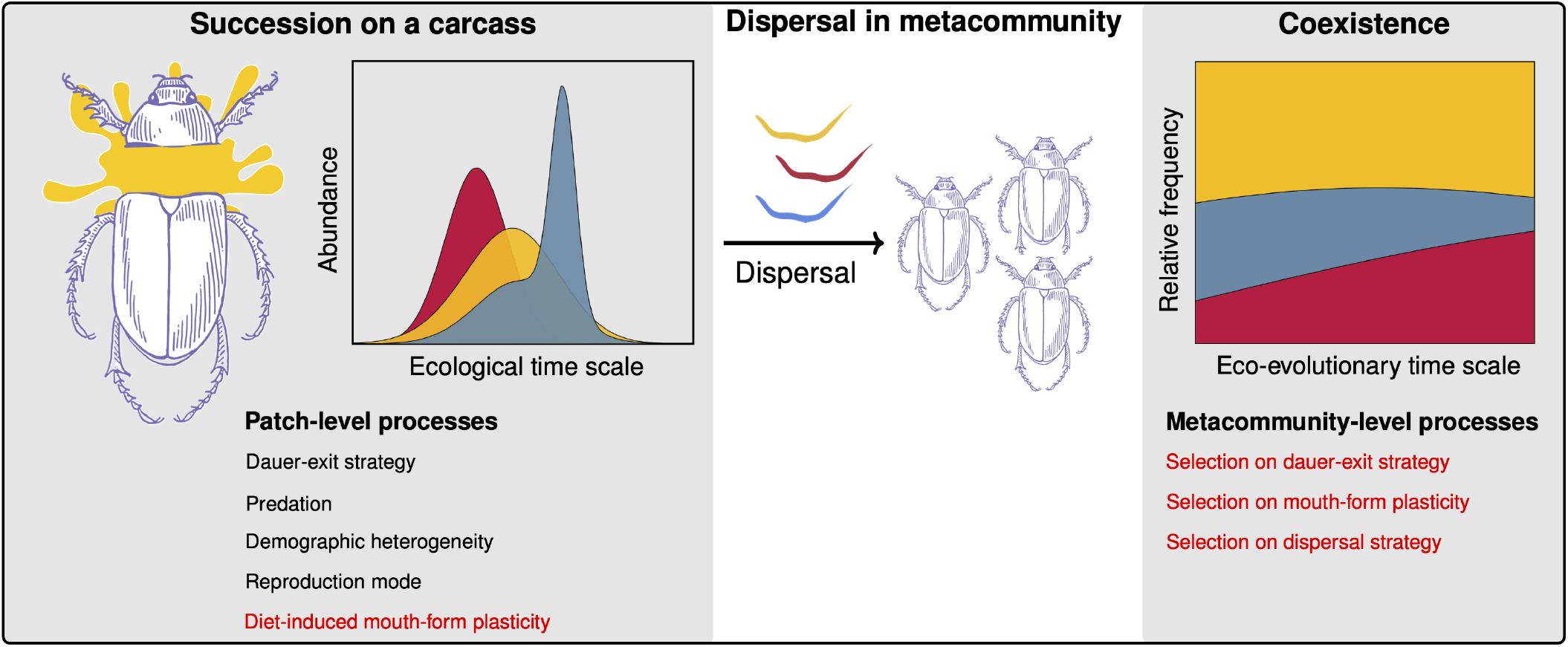
The role of succession and dispersal in coexistence on a decaying beetle as a model for metacommunity ecology. The decaying beetle can be envisioned as a transient patch, in which processes such as predation and strategies including dauer-exit timing and reproductive mode result in a specific pattern of succession. The dispersal at the metacommunity level can result in coexistence, given the constant dispersal and succession events at the metacommunity level. Additionally, selection could shape various traits of nematodes at the eco-evolutionary time scale to maximize their long-term fitness.

Despite the insights gained from our integrative study, crucial aspects remain unexplored. For example, the diversity of feeding structures in various *Pristionchus* species is a case of developmental (phenotypic) plasticity, where the population-level proportion of predatory to bacteriovorus nematodes is determined by various environmental factors (*47–50*). Most notably, the microbial diet strongly influences mouth-form development and can even result in transgenerational epigenetic inheritance (*51, 52*). We have previously explored how phenotypic plasticity can affect ecological competition (*31, 53*). However, it remains to be seen how the composition of the microbial community sampled from the decaying beetle – instead of the standard laboratory diet – would affect life-history traits. More importantly, the extent to which these natural diets would result in diet-induced plasticity in feeding structures is currently unknown and awaits future analysis that will profit from strong modeling contributions. In conclusion, the experimental and theoretical study of the decaying *G. buphthalma* beetle can be used to address various metacommunity and succession questions, resulting in a better understanding of long-term as well short-term coexistence in such systems – by combining empirical manipulations with agent-based modeling.

## Acknowledgments

We used the Opuntia cluster from the Hewlett Packard Enterprise Data Science Institute at the University of Houston. We thank the support from the Research Computing Data Core at the University of Houston.

## Authors contributions

RJS and AK conceived the project. PT observed the succession on beetle carcasses. PT and AK measured the life-history traits. AK wrote the code and visualized the results. RJS and AK wrote the manuscript.

## Data and materials availability

The software used to run all simulations and conduct all the data analysis was written in Python 3.11.7 with NumPy 1.26.4 (*54*). The Bayesian statistical analyses were done using PyMC 5.21.1 (*55*). The code and the raw experimental data are available at https://github.com/Kalirad/Communotron.

