## Supplementary Materials for "The dynamics of coexistence and succession in a decaying ecosystem"

### Materials and Methods

#### The structure of *Communotron*

Below, we describe our agent-based model using the Overview, Design concepts and Details (ODD) protocol [*1*].

##### Purpose

*Communotron* is designed to enable a quasi-realistic simulation of individual worms, as autonomous entities, that follow a life cycle, can interact with other entities in the environment and whose characteristics are influenced by the environment. These entities move on a 2-dimensional grid, a simulacrum of a decaying beetle and its immediate surrounding. Specifically, *Communotron* was designed to enable formation and decline of a community in order to study the boom-and-bust cycle observed in free-living nematodes that are found in association with scarab beetles. The model aims to capture the dynamics of these interactions and their ecological implications, providing insights into the community assembly processes.

##### Entities, state variables, and scales

In *Communotron*, each worm is simulated as an independent entity which, depending on its developmental stage, can move on a 2-dimensional grid. Each worm has a designated universally unique identifier. The probability of transition from developmental state $i$ to state $j$ is

$$P_{i \to j} = \frac{1}{1 + e^{-k(\tau - \tau_0)}} \quad , \tag{S1}$$

where $\tau$ is the current age of the worm and $\tau_0$ is the expected age at which the transition occurs. The temporal unit of the model is hour. During the life-cycle of a worm, alternative developmental stages are reached as a function of the environment. For egg $i$ at ($x_i$, $y_i$), normal development results in a juvenile. However, if at ($x_i$, $y_i$) the amount of resource available $< R_{\text{lim}}$, egg $i$ develops into an arrested juvenile. For juvenile $i$ at ($x_i$, $y_i$), if all the positions within a Moore neighborhood with range $r = 1$ around ($x_i$, $y_i$) are occupied, juvenile $i$ develops into a dauer instead of an adult. Both an arrested juvenile and a dauer larvae can re-enter the normal life cycle if resource availability in their location is $\geq R_{\text{lim}}$ and a Moore neighborhood with range $r = 1$ around their current position is not fully occupied (Note: eggs or dead worms are not counted when calculating occupancy in a Moore neighborhood).

Within radius $r$ around the center of the lattice, resource is available at $R_{\text{max}}(t)$, which changes during a simulation following

$$R_{\text{max}}(t) = \frac{1 + \text{Sin}(\omega \Delta t)}{2} \quad , \tag{S2}$$

where $t$ is the current time of simulation, $\Delta t = t - t0$, where $t0$ determines how long from the start of the simulation bacterial growth would start, and $\omega = 2\pi n_{\text{cycle}}$, enabling modification of the number of growth phases of the bacterial resource during the simulation. Parameter $n_{\text{cycle}}$ specifies the number of resource availability phases during the simulation.

The resource level at any position outside radius $r$ around the center– reflecting the growth and decline of bacterial load on the decaying carcass at the given point in time and space – follows

$$R_(x_i, y_i) = e^{-\delta(d-r)} \times R_{\text{max}}(t) \quad , \tag{S3}$$

where $\delta$ is the decline rate, $d$ is the Euclidean distance between the center and ($x_i$,$y_i$), and $r$ is the radius.

The gradient of bacterial resource in the environment affects the movement of worms. Juvenile and adult worms move in a biased random walk in the direction of the gradient. In contrast, dauer larvae move randomly, but biased against the gradient. Table S1 provides a summary of the parameters used in *Communotron*.

**Process overview and scheduling**

During each step of a simulation, the following steps are taken:

1. Motile stages – juvenile, adult, and dauer – move in a biased random walk on the grid. Predation occurs during this step, where *P. pacificus* adult $i$ can kill juveniles, arrested juveniles, or dauer larvae of the other species with probability $\psi$. The probability is a function of the age of the predatory adult:

$$\psi(\tau_i) = \eta e^{-\theta\tau_i} \quad , \tag{S4}$$

   resulting in a decline in the ability to engage in surplus killing due to aging.

2. The adults lay their eggs in a Moore neighborhood with range $r = 1$ around their current position. The number of eggs laid per hour is based on the mean number of viable progeny per day for each nematode species. To use these values, the daily mean values are uniformly distributed for each worm over $24$ hours.

3. The decision to develop to the next developmental stage is taken by comparing a random number sampled from a uniform distribution with $P_{i \to j}$ (Eq. S1). The alternative developmental paths are taken as specified above. The feeding structures are also updated in cases where an adult is developed from a juvenile or dauer larva.

4. Survival of adults, arrested juveniles, and dauer larvae are calculated based on a sigmoidal transition function identical to Eq. S1.

5. The simulation clock moves an hour forward.

**Design concepts**

*Basic principles:* The purpose of *Communotron* was to enable an agent-based framework with quasi-realistic life-cycles, the ability to include experimental data, resource dynamics, and simulation on the scale of hours. Specifically, *Communotron* addresses a number of the limitations of our previous agent-based framework [*2*], namely the absence of resource dynamics and restricted spatial components. Additionally, the emigration of dauer larvae in *Communotron* is tied to the biased random walk of worms in this stage.

*Emergence:* The probabilistic nature of interactions between the worms compounded by the effect of the explicit spatial aspect of the model results in emergent dynamics that deviate from a deterministic model.

*Learning:* The model does not include any learning.

*Prediction:* *Communotron* is designed to predict how the emergence of nematodes on a decaying beetle, coupled with their feeding structure, their modes of reproduction, and their fecundities, shapes the community composition during the decomposition of the beetle carcass.

*Interaction:* In *Communotron*, predation of juveniles, arrested juveniles, and dauer larvae of competing species by *P. pacificus* adults is the direct interaction between the agents. By affecting the local densities, species indirectly affect each other by influencing the likelihood of transitioning to alternative developmental stages.

*Observation:* Each worm in *Communotron* has a designated universally unique identifier, enabling tracking the worms during the simulations. The age, developmental stage, location, migration status of worms are recorded during the simulation, as well as killing events.

**Initialization**

The parameters used to start a simulation are specified in Table S1. Additional information concerning the initial parameters can be found in figure legends. The initial number of each nematode species is drawn from a Poisson distribution based on the specified expected numbers provided by the user.

#### Laboratory Assays

##### Community on a plate

*G. buphthalma* beetles sampled from Réunion island during a sampling trip in January 2024 were used to construct "community on a plate", where each sacrificed beetle is placed on a nematode growth medium (NGM) plate. Following the outgrowth of bacteria and fungi from the carcass, nematodes in their dauer state emerge from the carcass and develop into adults. The plates were observed in regular intervals for 120 hours following the sacrifice of the beetle in order to record the timing and the identity of the nematodes emerging from the carcass. The nematode species were identified morphologically based on the published descriptions of *P. pacificus*, *P. mayeri*, and *Acrostichus* [*3–5*]. Subsequently, the identity of the nematodes were confirmed genetically using the following 18S primers: SSU988F (5'-CTCAAAGATTAAGCCATGC-3') and SSU2646R (5'-GCTACCTTGTTACGACTTTT-3').

##### Fecundity

To measure the fecundity of the isolates of *P. pacificus* and *P. mayeri*, both self-fertilizing hermaphrodites, we transferred single worms to fresh plates – seeded with 20 $\mu L$ *Escherichia coli* OP50 – every 24 hours for 7 days. For each day, viable juveniles observed on plates after $48$ hours were recorded. For each isolate of *P. pacificus*, $40$ hermaphrodites were transferred daily ($120$ mothers in total). In the case of *P. mayeri*, $60$ hermaphrodites of each strain ($180$ mothers in total) during a $7$ day period. For both species, given the low number of viable progeny in the last three days of the assay, the progeny for days $5-7$ were grouped together. For the gonochorist *Acrostichus* sp. RSO013, a pair of male and female – both at J4 larval stage – were transferred every 24 hours to a freshly-seeded plate with *E. coli* OP50 for 17 days. We transferred $30$ pairs daily. During the course of the experiment unhealthy or dead males was replaced. Given the missing information on fecundities on days 14 and 16, we used the mean fecundities of day $i+1$ and day $i-1$ for these two days, where $i \in \{14, 16\}$.

##### Sex ratio

The sex ratio of *Acrostichus* sp. RSO013 was estimated by counting the number of male and female progenies during the fecundity assay of this isolate.

##### Feeding structures

Three isolates of *P. pacificus* and three isolates of *P. mayeri* were grown on *E. coli* OP50 under standard laboratory conditions and $750$ adult worms from each isolate ($4500$ worms in total) were examined morphologically – based on the mouth width and the shape of the dorsal tooth.

#### Statistical analyses

##### Fecundity parameter

To estimate difference in the total number of progeny between species of *Pristionchus* and *Acrostichus*, we constructed a bayesian model:

$$\begin{aligned} y_i &\sim \mathrm{Poisson}(\lambda_j) \quad , \\ \lambda_j &\sim \mathrm{Gamma}(\alpha, \beta) \quad , \end{aligned} \tag{S5}$$

where $y_i$ is the number of viable progenies of individual $i$ of isolate $j$. For the gamma prior, $\alpha = 2$ and $\beta = 1$. The model was fitted to the laboratory measurements using PyMC [*6*] to estimate $\lambda$ for each isolate. We ensured that the estimated $\lambda$ values were stable using common convergence diagnostics with effective sample sizes $\geq 10,000$ [*7*].

### Supplementary Tables

| Parameter | Description | Value |
|---|---|---|
| $m \times m$ | Determines the size of the grid | $100 \times 100$ |
| $r$ | The radius of the resource | $20$ |
| $\delta$ | The decline rate of the resource using in Eq. S3 | $0.3$ |
| $k$ | Determines the shape of Eq. S1 | $0.1$ |
| $R_{\text{lim}}$ | The lower limit of resource level used for alternative developmental paths | $0.4$ |
| $\eta$ | Predation parameter used in Eq. S4 | $0.005$ |
| $\theta$ | Predation parameter used in Eq. S4 | $0.01$ |
| $L$ | Number of steps taken during random walk by each agent at each step | $10$ |
| $\tau_0$ (Egg) | The parameter used in Eq. S1 to calculate transition probability form egg to juvenile or arrested juvenile | $48$ |
| $\tau_0$ (Juvenile) | The parameter used in Eq. S1 to calculate transition probability form juvenile or adult or dauer | $72$★, $96$☆, $120$† |
| $\tau_0$ (Dauer) | The parameter used in Eq. S1 to calculate dauer exit into adult | $72$★, $96$☆, $24$† |
| $\sigma_0$ | The parameter used to calculate survival probability | $500$ (A), $1000$ (D, AJ) |

Table S1: **The main parameters used in *Communotron.*** ★: *P. pacificus*; ☆: *P. mayeri*; †: *Acrostichus* sp.; A: adult; D: dauer; AJ: arrested juvenile.

#### Supplementary Figures

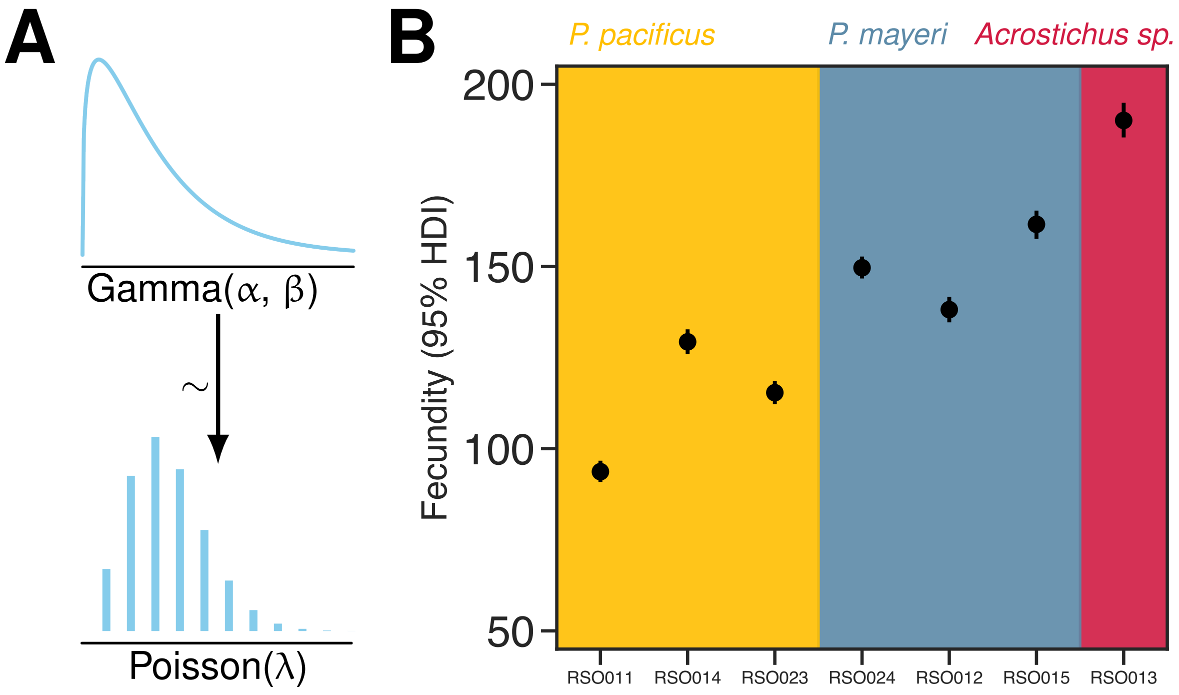


Figure S1: **(A)** A Kruschke-style model diagram, showing the structure of the bayeisan model used to estimate the life-time fecundity of the nematodes based on our experimental measurements. The model assumed that for each isolate, the total number of viable progenies can be explained as a Poisson process with an isolate-specific fecundity parameter ($\lambda$). (B) The estimated $95\%$ highest density interval (HDI) for the fecundity of isolates of three nematode species.

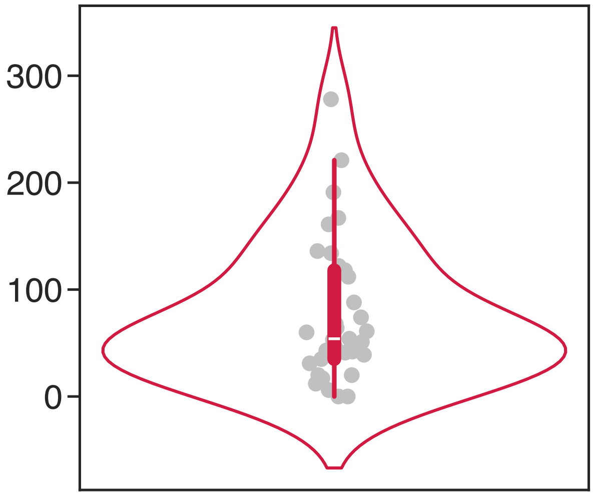


Figure S2: The observed number of nematodes on the beetle carcasses $24$ hours following the sacrifice of the beetle. All the nematodes emerged at this point belonged to *Acrostichus* sp.
